# Rapid and label-free exosome analysis by surface-enhanced Raman spectroscopy using silver nanoparticle substrate based on selective laser ablation and melting of silver nanowire

**DOI:** 10.1101/2024.01.13.575493

**Authors:** Hyeono Nam, Jong-Eun Park, June Sik Hwang, Seunggyu Kim, Seong Jae Kim, Sanha Kim, Minyang Yang, Jessie S. Jeon

## Abstract

Early diagnostics of breast cancer is crucial to reduce the risk of cancer metastasis and late relapse. Exosome, which contains distinct information of its origin, can be the target object as a liquid biopsy. However, its low sensitivity and inadequate diagnostic tools interfere with the point-of-care testing (POCT) of the exosome. Recently, Surface-enhanced Raman spectroscopy (SERS), which enables the detection of Raman scattering, has been proved as a promising tool for exosome detection but the fabrication process of SERS probe or substrate is still inefficient and far from large-scale production. This study proposes rapid and label-free detection of breast cancer-derived exosomes by statistical analysis of Raman spectra using silver nanoparticle SERS substrate fabricated by selective laser ablation and melting. Employing silver nanowire and optimizing pulse repetition rate power enable rapid and energy-efficient fabrication of SERS substrate. The functionality including sensitivity, reproducibility, stability, and renewability was evaluated using rhodamine 6G as a probe molecule. Then, the feasibility of POCT was examined by the statistical analysis of Raman spectra of exosomes from malignant breast cancer cells and non-tumorigenic breast epithelial cells. The presented framework is anticipated to be utilized in other biomedical applications, facilitating cost-effective and large-scale production performance.

## 1. Introduction

Breast cancer, one of the most fatal diseases, is significantly increasing its mortality rate and has high metastatic potential to other organs as well as a high probability of late relapse after treatment [1–3]. To decrease the level of minimal residual disease, the importance of developing effective treatment and early-stage diagnostics cannot be overstated [3]. Exosome, which is an extracellular vesicle encapsulated by a phospholipid bilayer, is generated in multivesicular bodies and released through the process of exocytosis [4–7]. Hence, this nanoscale structure includes cellular information from which it is generated [8]. Since exosomes exist almost everywhere in the body including plasma [9, 10], urine [11, 12], breast milk [10, 13], and saliva [10, 14], utilizing these exosome-containing body fluids as a liquid biopsy takes advantage of non-invasiveness, contamination-free, early diagnosis, and reducing the burden on the patient [15, 16]. Despite these clinical potentials, diagnosis by detection and evaluation of the exosomes is difficult to apply to actual clinical fields due to the lack of sufficient and high-sensitivity equipment to detect them.

Surface-enhanced Raman spectroscopy (SERS), which is a highly sensitive detection technique based on the amplification of Raman scattering in the presence of the metallic nanostructure interface, has been widely exploited to identify Raman scattering of target molecules otherwise difficult to detect due to its weak signal intensity [17, 18]. Statistical analysis of spectra derived from molecular structure enables to perceive not only various constituents of a substance, but also the presence of molecules with a small composition ratio. In recent years, nanostructures to implement SERS engage with body fluids, realizing SERS as one of the liquid biopsy tools to replace traditional tissue biopsy [19]. Most of all, SERS-based characterizations of cancer-derived exosomes have been extensively investigated. Labeling methods that allow SERS probes to naturally bind to individual exosomes through principles such as electrostatic adsorption [20–22], hydrophobic interaction [23, 24], and aptamer or peptide-based selective binding [25, 26] have the advantage of obtaining noise-reduced spectra from individual exosomes thus simplify the identification of the origin in the mixture. Nonetheless, chemical reaction-based SERS probe manufacturing process is complicated, expensive and requires a significant amount of time and labor. On the other hand, label-free methods that samples are merely deposited on a nanopatterned substrate have the advantages of a simplified process by reduced substrate preparation time with providing a large amount of exosomal data efficiently compared to the previous methods [27–31]. Nevertheless, current SERS substrate fabrication processes are still complicated, time-consuming, laborious, and limited for large-scale production to apply point-of-care testing (POCT).

Laser ablation borrowing the power of inducing the formation of high-temperature, high-density plasma plume on the surface of the material facilitates fabrication of nanoparticles as well as enables nanopatterning in a deterministic manner depending on the properties of material and laser conditions [32, 33]. This top-down nanofabrication approach has given assistance to produce monodisperse colloidal nanoparticle solutions [34–40], synthesized nanoparticles through carrier gas [41–44] and nanopatterned surfaces [45–47] by employing a wide range of materials. While this method still has the hurdle of requiring high input energy to deal with bulk materials that burden with expensiveness and inefficiency [48], the potential of mass production in a facility with rectified shortcomings will make laser ablation-based SERS substrate fabrication take a step closer to POCT.

This study presents rapid and label-free detection of breast cancer by statistical analysis of Raman spectra of the breast cancer-derived exosomes using silver nanoparticle (AgNP) SERS substrate. To overcome the abovementioned current hindrances, selective laser ablation and melting using a nanosecond-pulse laser was coupled with silver nanowire (AgNW), which has the commercial potential of large-scale and low-cost production [49], to fabricate AgNP SERS substrate. Furthermore, optimization of pulse repetition rate and power allowed highly compacted, uniform, and single-layered AgNPs and permitted efficient input energy operation. Using rhodamine 6G (R6G) as a probe molecule, the fabricated substrate was confirmed its functionality and capability by quantitatively analyzing performance abilities such as sensitivity, reproducibility, stability, and renewability. Lastly, statistical analysis of the spectra from three malignant breast epithelial cancer cell-derived exosomes and non-tumorigenic breast epithelial cell-derived exosomes was carried out to prove the applicability of the substrate as POCT tools. The presented fabrication method and SERS platform, which reduce the burdensome preparation time and cost and do not require labeling for the detection, is expected to be applied to various POCT platforms to encourage the biomedical field.

## 2. Results and Discussion

### 2.1. AgNP substrate optimization

Figure 1a illustrates the overall process of this study. Vacuum filtration of AgNW solution makes AgNWs adhere to the glass substrate with nearly monolayer. By nanosecond pulse laser ablation, two distinctive areas are appeared by plasma expansion; vaporized zone and melted zone (Figure 1b). While relatively small AgNPs comprise in the vaporized zone, melted zone with lower temperature enables to Ag vapor plume to coagulate and consequently generate larger AgNPs to some extent (Figure 1c). Combining with the AgNW quasi-monolayer and selective laser ablation and melting, a highly monodispersed AgNP substrate was fabricated (Figure 1d).

**Figure 1.**
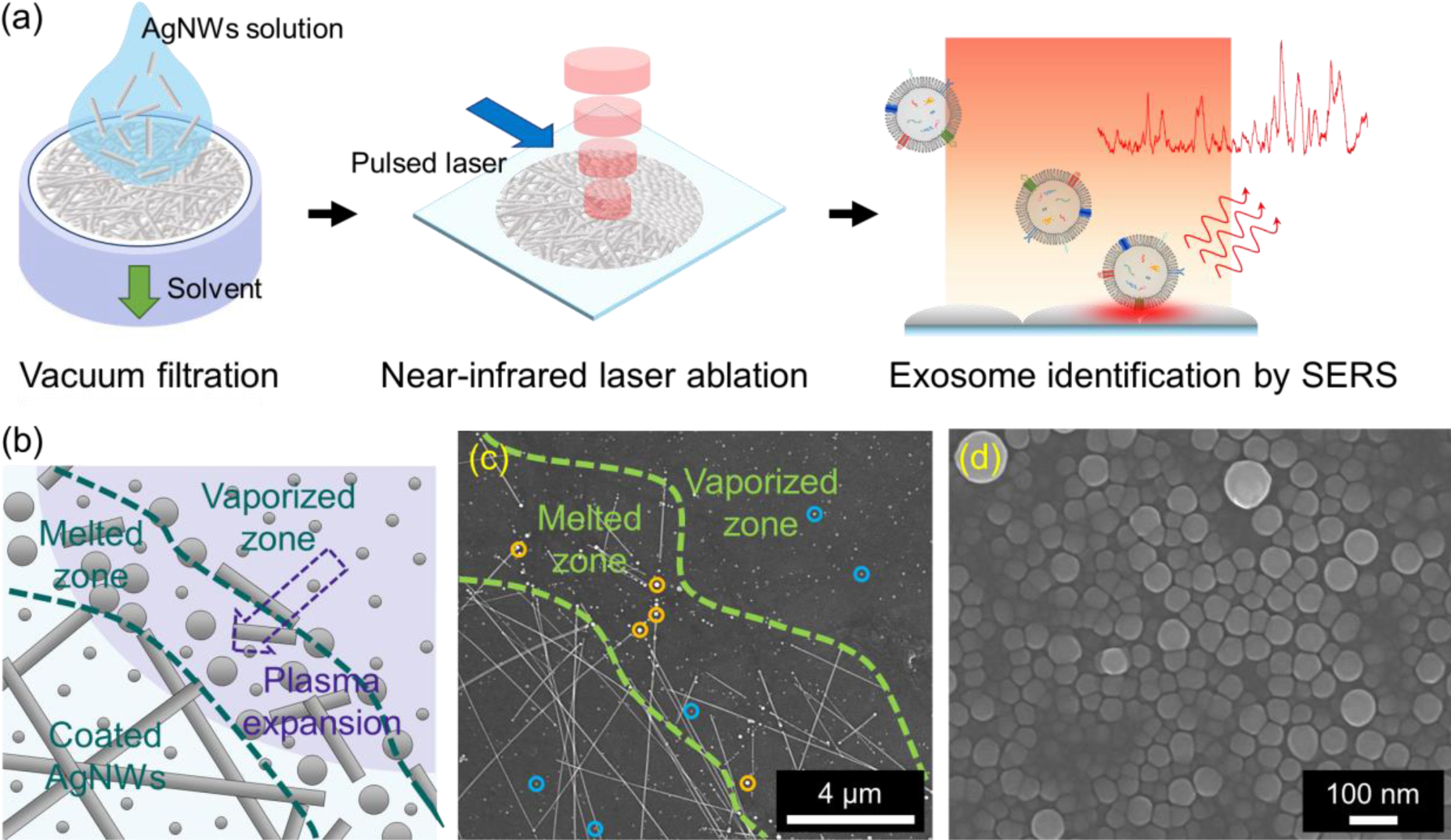
(a) Schematic of the research overview. (b) AgNPs are developed by direct melting (big AgNPs) and plasma expansion (small AgNPs). (c) SEM image at boundary between coated AgNWs and developed AgNPs. (d) Magnified SEM image of laser ablated region displays highly compacted AgNPs.

Optimization of manufacturing conditions (AgNW concentration, laser power, pulse repetition rate, ablation time) was implemented for cost-effective, energy-efficient fabrication as SERS substrate (Figure 2, 3; Figure S4, Supporting Information). To quantify the effect of the AgNW concentration on nanoparticle size distribution, two AgNW solutions with different concentrations were employed (Figure 2a-f). As shown in Figure 2c, f, solution with high concentration offers a more homogenous size distribution, as well as a bigger mean nanoparticle size, compare to low concentration. This phenomenon is presumed to be the influence of the melting zone that brought about plurality of larger nanoparticles by acquiring increased vaporized Ag source from AgNWs with high concentration.

**Figure 2.**
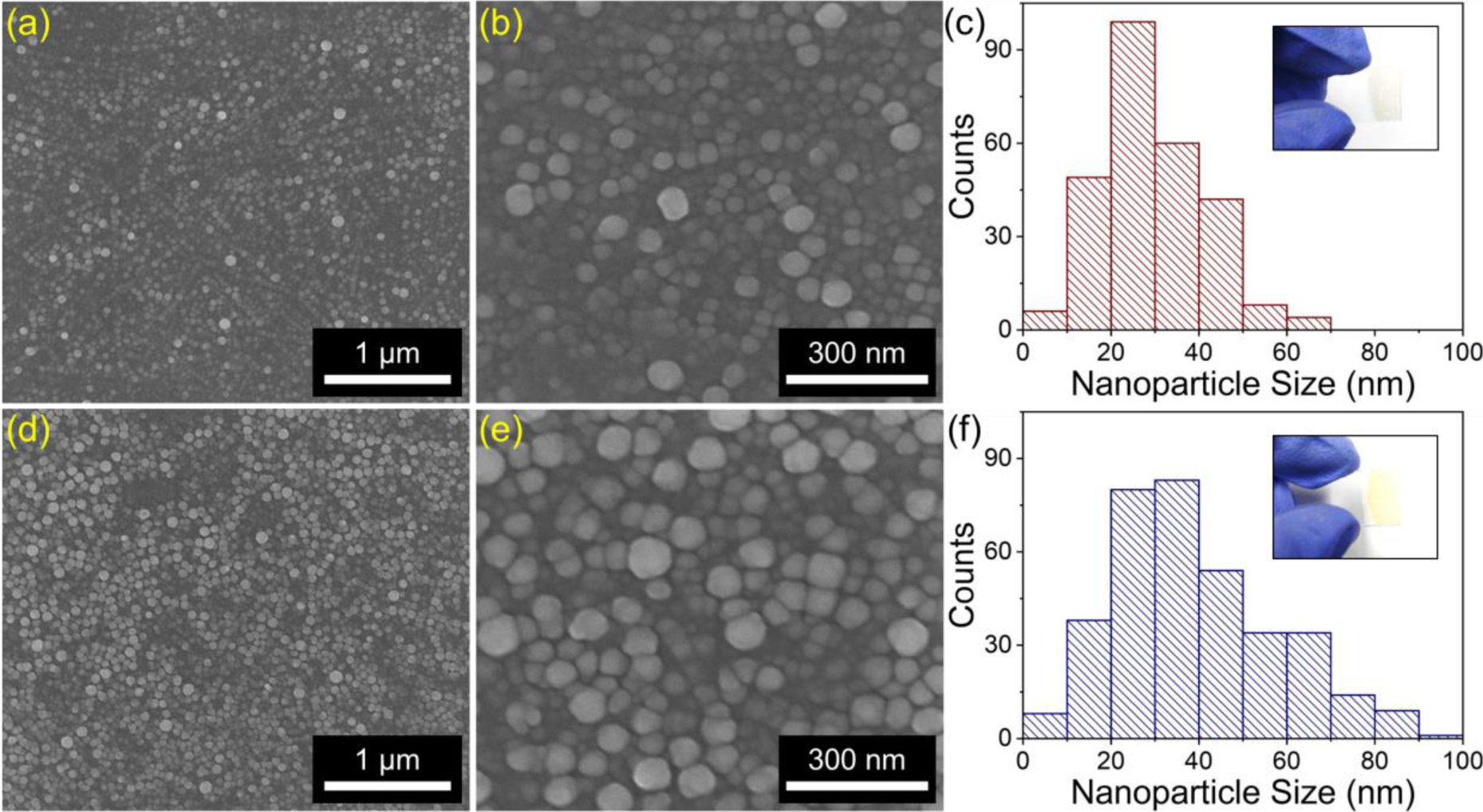
(a, b) SEM images of AgNPs from AgNWs. (c) Size distribution of AgNPs from AgNWs with low concentration. (d, e) SEM images of AgNPs from high concentration of AgNWs. (f) Size distribution of AgNPs from AgNWs with high concentration.

For large-scale production, high input energy is one of the abstruse hurdles to overcome [50]. By varying the power and pulse repetition rate, the minimum required input energy for contriving highly compact AgNPs was examined (Figure 3a-c). On the spin-coated AgNW substrate, laser ablations were carried out under 16 different conditions; 4 different powers by 4 different pulse repetition rates (Figure 3a). From each condition, scanning electron microscopy (SEM) images of AgNP surfaces were obtained and evaluated by quantifying the covered area and the count of AgNPs (Figure 3b, c). With a few exceptions, there is a conspicuous propensity that is hard to covert AgNWs to AgNPs under 0.02 mJ. This is speculated that there should be more than 0.02 mJ for vaporization and melting to deal with AgNWs in our framework. For laser energy with 0.04 mJ, 0.03 mJ, and 0.027 mJ, significant increase in the number of AgNPs and covered area (particle number of 174, 113, and 190; covered area of 5.27 %, 2.5 %, and 5.71 % for each condition) were observed. Following these conditions, the number of AgNPs and covered area were increased successively as the laser energy increased (173 and 4.12 % at 0.06 mJ; 241 and 5.74 % at 0.04 mJ). Laser condition of 20 W and 250 kHz (0.08 mJ) rendered 393 AgNPs with 9.24 % of covered area in the quadrat of SEM image. With the same pulse repetition rate, vacuum filtered AgNW substrate represented the similar tendency that the increased laser power contributed more compact and smaller AgNPs which assigned amplified Raman spectrum of R6G (Figure S4a, c, Supporting Information).

**Figure 3.**
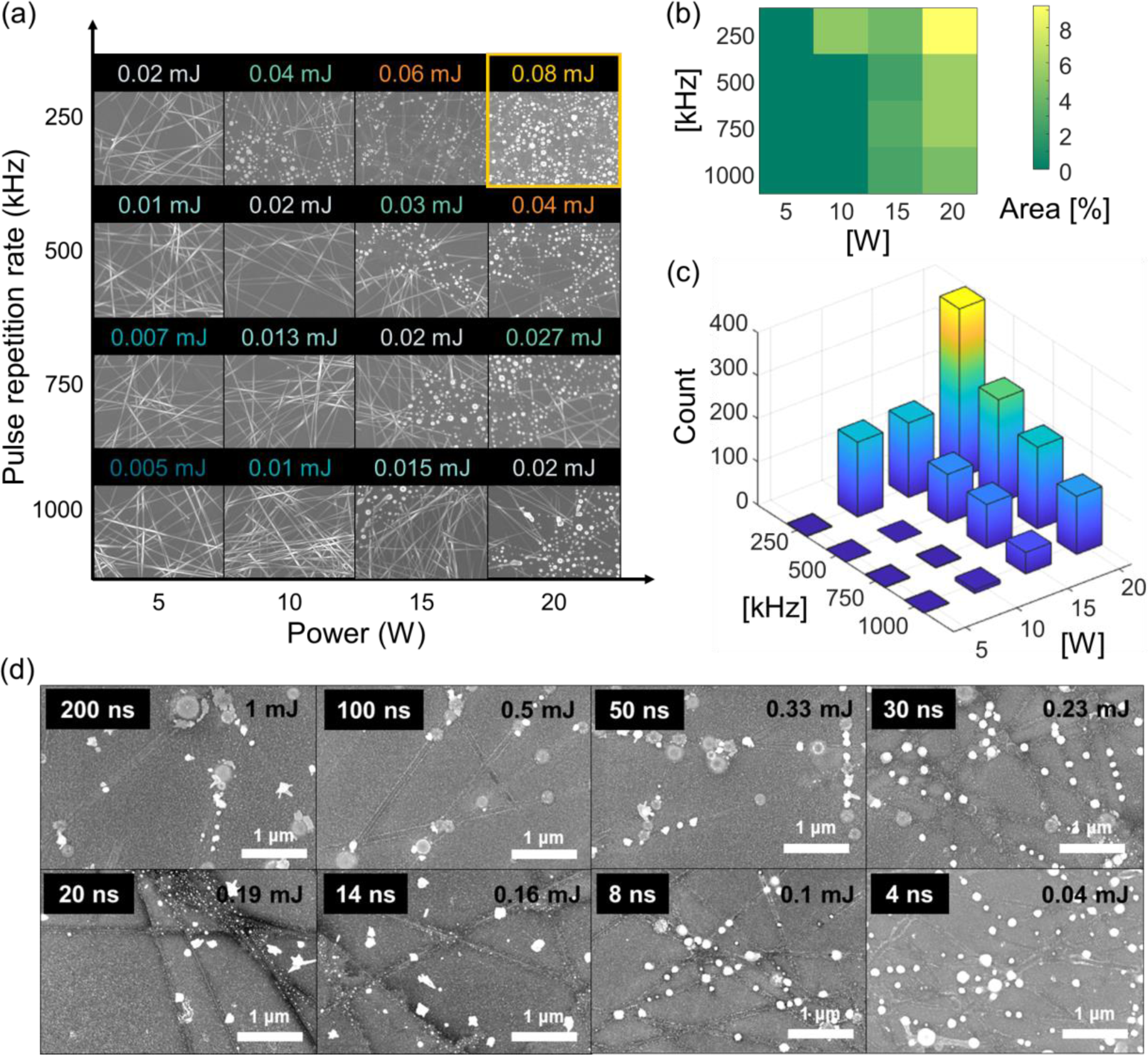
(a) SEM images of laser ablated AgNW substrate with respect to laser power and pulse repetition rate. (b) Heat map of the percentage of covered area by AgNPs to visualize the efficiency according to laser conditions. (c) Number of AgNPs on the substrate with regard to each condition. (d) SEM images of laser ablated substrate under the fixed average power with maximum input energy.

We further tested AgNP development under the fixed average power with maximum input energy (Figure 3d). Prolonging the duration time could increase the total input energy under the same average power. In contrast with the previous results (Figure 3a-c), excessive input energy, rather than moderate one, hampered the formation of AgNPs (Figure 3d). We postulated that comparatively enhanced vaporization impedes ordinary condensation of sublimated AgNW vapor so that incurs non-circular, smeared Ag patterns on the substrate. In the same context, immoderate ablation repetition time revealed AgNP substrate with distorted particle shape and reduced amplification of R6G Raman spectrum (Figure S4b, d, Supporting Information). Taken together, sufficient but not excessive input energy was established in our framework for AgNP substrate fabrication with a facile and affordable process.

### 2.2. AgNP substrate performance characterization

To make use of AgNP substrate at SERS, functional properties such as sensitivity, reproducibility, stability, and renewability should be evaluated and validated [51]. R6G, which expresses distinctive peaks by resonance Raman scattering, was engaged as a probe molecule [52]. The enhancement factor (*EF*) calculated before sensitivity measurement enables to estimate our substrate performance; (*EF* = (*Isers***×***Nnr)*/(*Inr***×***Nsers)*), where (*Isers*) and (*Inr*) are the SERS and normal Raman intensities, respectively; (*Nsers*) and (*Nnr*) are the number of probed molecules by SERS and normal Raman, respectively. Sensitivity was assessed by decreasing the concentration of R6G by a factor of ten for each step (Figure 4a). From 10 nM to 10 pM, almost characteristic peaks of R6G faded except a few prominent peaks (612 cm^−1^, 773 cm^−1^, 1362 cm^−1^, and 1650 cm^−1^). Among these peaks, the magnitude of the signal at 1650 cm^−1^ was hired to quantitatively investigate the limit of detection (LOD) (Figure 4b). In the logarithmically scaled coordinate, the Raman intensities for each concentration were fitted using linear regression with an r-squared (*R^2^*) value of 0.98 and LOD was calculated.

**Figure 4.**
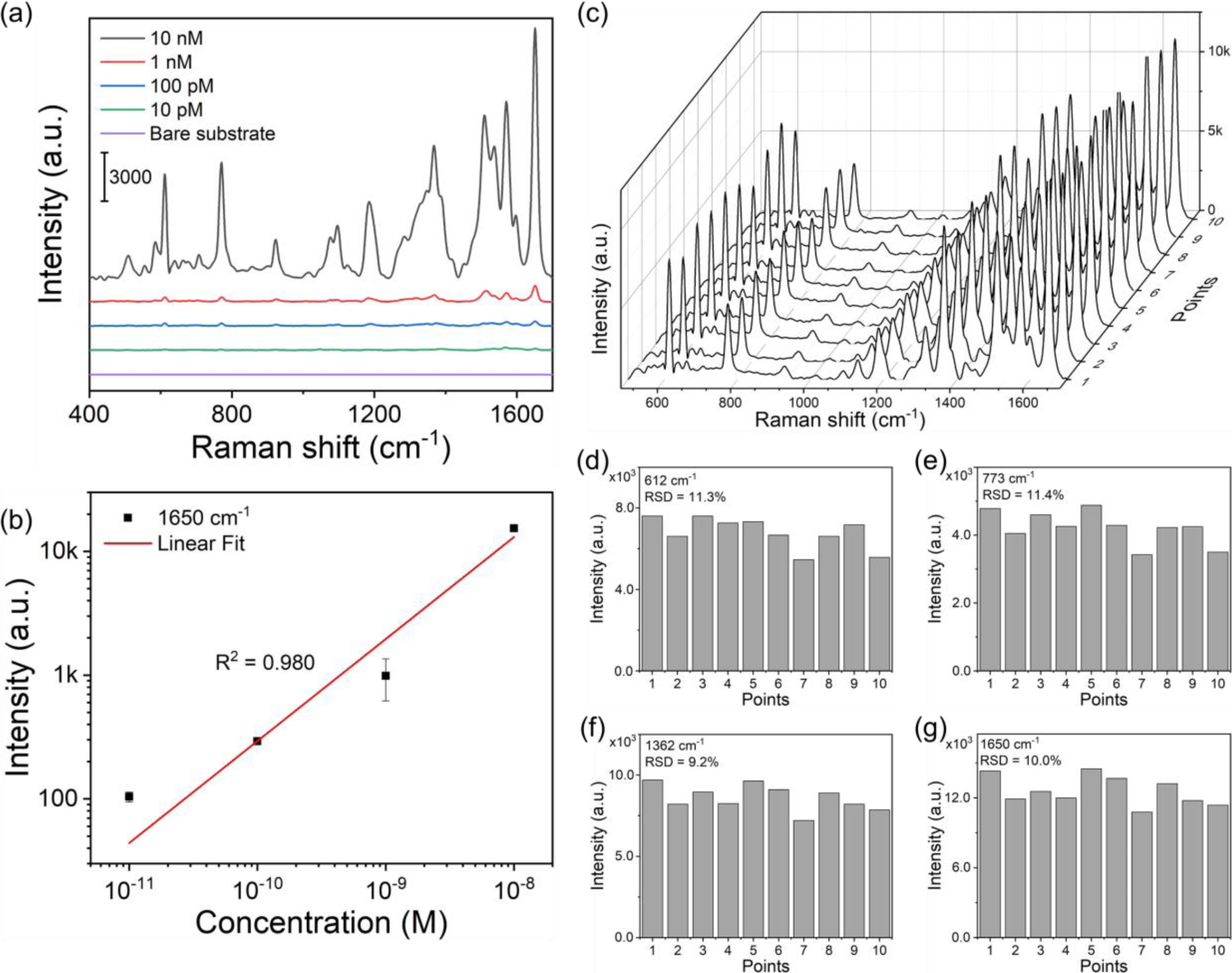
(a) Raman spectra of R6G with different concentrations. (b) Signal intensities at 1650 cm^−1^ with regard to R6G concentration for the sensitivity evaluation (error bar: standard deviation). (c) Raman spectra of R6G from ten randomly selected locations. (d) Signal intensities comparisons of four distinct wavenumbers.

The reproducibility takes the portion of importance in terms of collecting a large amount of data with accuracy [51]. Followed by confirming the even surface morphology of AgNPs by Atomic Force Microscopy (AFM) (Figure S2d, Supporting Information), specific peaks (612 cm^−1^, 773 cm^−1^, 1362 cm^−1^, and 1650 cm^−1^) from 10 randomly obtained Raman spectra were compared to appraise reproducibility (Figure 4c-g). Figure 4c represents the entire profiles of 10 randomly chosen Raman spectra individually. As we perceive that there is more probability to entail high deviation at outstanding intensities by intuition, a comparative analysis was conducted for the specific peaks (Figure 4d-g). Four specific peaks (612 cm^−1^, 773 cm^−1^, 1362 cm^−1^, and 1650 cm^−1^) have relative standard deviation (RSD) of 11.3 %, 11.4 %, 9.2 %, and 10.0 %, respectively. We postulate that somewhat but not significant RSD values might be derived from the relatively uneven district by sparsely existing sizable AgNPs.

As high temperature incurs oxide films on the surface of the silver [53], we confirmed the oxidation state of the AgNPs by transmission electron microscopy (TEM) (Figure S2 a-c, Supporting Information). Figure S2b, c denote the atomic distribution of silver and oxygen, respectively, which implies the formation of oxide film covering the AgNPs. The result from time-of-flight secondary ion mass spectrometry also indicates the existence of oxide film by the ratio of silver oxide peak and silver peak (Figure S3, Supporting Information). Although the formation of oxide film protects from the oxidation penetrating the AgNPs, oxidation itself could affect the performance [54]. The stability of the substrate was evaluated by characterizing four specific peaks (612 cm^−1^, 773 cm^−1^, 1362 cm^−1^, and 1650 cm^−1^) of R6G every week (Figure S5a-e, Supporting Information). Except for the 773 cm^−1^ and 1362 cm^−1^ peaks at the first week, all of the signals diminished gradually. Since the natural oxide thickness on the silver converges [55], we assume the reason for decreasing signal intensity from crumbled regions. Renewability was assessed by the comparison of two Raman spectra from two independent sample loadings on the same substrate (Figure S5f-j, Supporting Information). Alike the stability measurement, the intensities of four different peaks from R6G were compared respectively. All the intensities at the second sample loading recovered almost as much as those at the first sample loading. Nevertheless, slight but not significant decreases in signal intensities were identified that are presumed to be the same reason in the stability test. Because the AgNWs are merely attached to the glass substrate by vacuum filtration, we expect to improve the stability and renewability through enhancing the AgNWs adhesion to the substrate by utilizing another adhesional method such as roll-to-roll technique [56].

### 2.3. Statistical analysis of exosome-derived spectra: proof-of-concept

Exosomes from four different cell lines were used to investigate the prospect of our substrate in POCT (Figure 5). Exosomes from three malignant breast cancer cell lines and one non-tumorigenic breast epithelial cell line were isolated by differential ultracentrifugation [57]. The size distribution and concentration of each exosome suspension were analyzed by nanoparticle tracking analysis (NTA) (Figure S6, Supporting Information). Mode size and concentration for each exosome suspension were as follows: 132.0 nm and 1.28**×**10^11^ particles/ml for MCF7 exosome suspension, 133.2 nm and 2.09**×**10^11^ particles/ml for MCF 10A exosome suspension, 130.8 nm and 3.66**×**10^11^ particles/ml for SK-BR-3 exosome suspension, and 126.2 nm and 1.54**×**10^11^ particles/ml for MDA-MB-231 exosome suspension. Additionally, we confirmed the existence of exosomes by TEM and Western blot (Figure S7, Supporting Information). Figure S7a-d represent exosomes from each cell line of a donut-like shape with a hollow in the middle. Tetraspanins, such as CD9, CD63, and CD81, are commonly used markers to identify exosomes that exist on the membrane of the exosomes [58]. Evident protein bands of CD9 and CD63 were stained from each exosome suspension (Figure S7e, Supporting Information). Furthermore, tumor susceptibility gen 101 (TSG101) and annexin A2 (ANXA2), which are abundant in exosomes [59, 60], were also identified with strong protein bands.

**Figure 5.**
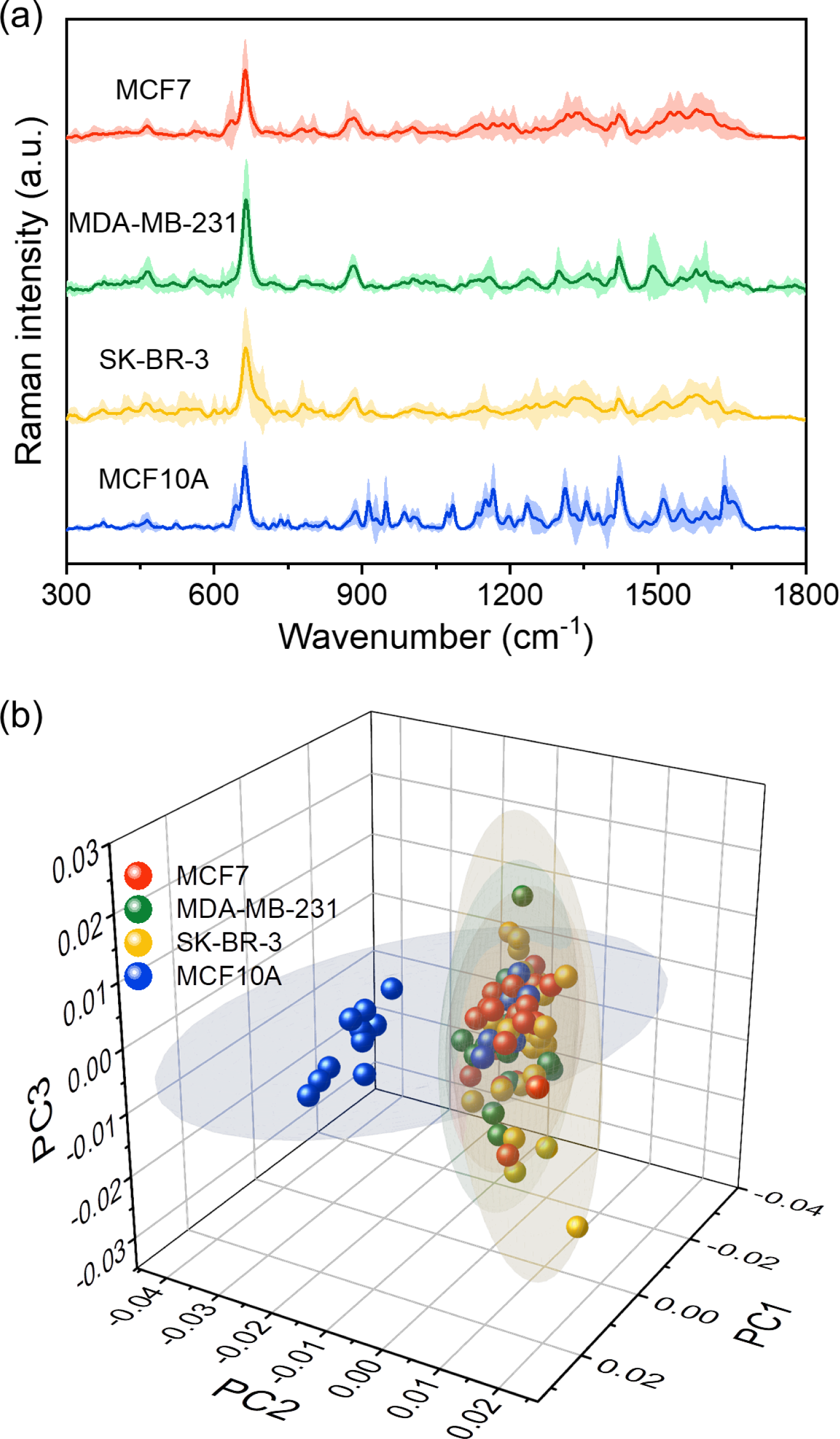
(a) Raman spectra from four different cell line-derived exosomes (shaded area: standard deviation). (b) PCA with three principal components reveals particular compartmentation of non-tumorigenic breast epithelial cell-derived exosome from malignant breast cancer cell-derived exosomes (shaded region: 95 % confidence ellipse).

Raman spectra with four disparate aspects were obtained from previously mentioned exosome suspensions using AgNP substrate (Figure 5a). Although some discrepancies can be recognized intuitively, the multidimensional attribute of Raman spectrum makes it hard to compare the spectra evidently. Principal component analysis (PCA) was performed to reduce the dimension of the Raman spectra and expedite the comparative analysis (Figure 5b). Three principal components (i.e. PC1, PC2, and PC3) were exploited based on scree plot which represents that the first three principal components explain nearly 35 % of the variance of the entire data (Figure S10b, Supporting Information). Except Raman spectra from MCF 10A-derived exosomes, the spectra originated from three different cell lines share their 95 % confidence ellipses (Figure 5b). Distinguishable categorization of malignant and non-tumorigenic groups by PCA not only verifies AgNP substrate as SERS platform but also implies the significant role of exosome as a clinical diagnosis. To inspect what wavenumbers substantially contribute to malignant/non-tumorigenic classification, loading plots were used to assess their weight on each principal component (Figure S9a, Figure S10a, Supporting Information). Although PC1 possesses the highest percentage of the total variance, PC2 manifests the characteristic that distinguishes non-tumorigenic breast epithelial cells from malignant breast cancer cells so that PC2 was preferentially chosen for the following comparison. By sorting out the specific wavenumbers, there are prominent propensities; peaks that have negative weight on PC2 are clear in non-tumorigenic breast epithelial cell-derived exosome, whereas peaks that have positive weight on PC2 are clear in malignant breast cancer cell-derived exosomes (Figure S9b, Supporting Information). We speculated that genetically different characteristics induced dissimilar cellular constituents which corresponds to each wavenumber (Table S1, Supporting Information).

## 3. Conclusion

In this study, selective laser ablation and melting methodology is introduced for rapid and label-free statistical analysis of exosomes by employing SERS. AgNW is utilized as a procurement material for AgNP SERS substrate. The concentration of AgNW and laser conditions (i.e., pulse repetition rate and power) are optimized for densely composed AgNPs with nano-scale cracks in which Raman scattering is amplified. This functional substrate has been proven for its versatility as SERS substrate by quantifying the sensitivity, uniformity, stability, and reproducibility. The subsequent statistical analysis of exosomes enables the distinction of malignant breast cancer cells from non-tumorigenic breast cells. While the abovementioned results confer potentiality on our substrate, there are several tasks to be improved for POCT and large-scale production. One of them is to skip the sample isolation or refinement process for user-friendly and time-saving POCT. Recently developed on-chip exosome isolation technologies could give inspirations to spur our platform for the integrated POCT system [61]. Additionally, the automated AgNW film fabrication such as the roll-to-roll technique can replace the vacuum filtration for large-scale production with high stability and portability [56]. As the current approach benefits from simplicity and diminishing significant time, we believe that the described methodology can establish a milestone for POCT of diverse diseases.

## 4. Experimental Section

### Scanning Electron Microscopy (SEM)

The observation region of the AgNP substrate was diced using a diamond cutter. The piece of AgNP substrate was fixed on the sample stand using conductive carbon tape. Colloidal silver paste (Ted Pella Inc, USA) was covered along the edges of the sample. The processed sample was coated with osmium and observed by field emission scanning electron microscope (Magellan400, FEI, USA) at 10 kV and 50 pA.

### AgNP Substrate Performance Test

Sensitivity was measured using different concentrations of R6G solutions. Sequential dilution with a factor of ten was implemented from 10 nM to 10 pM. To quantify the sensitivity, signal intensities at 1650 cm^−1^ was chosen and LOD were calculated. Raman spectra of R6G were obtained from ten randomly selected domains on the substrate to assess reproducibility. Signal intensities of four different peaks from obtained Raman spectra were used to calculate RSD. For the stability test, the fabricated substrates were kept in a dry, cool place until use. Every week, one of the stored substrates was used to measure the Raman spectrum of R6G. Signal intensities of four distinct peaks (612 cm^−1^, 773 cm^−1^, 1362 cm^−1^, and 1650 cm^−1^) from each week were compared. For the renewability measurement, the substrate on which Raman spectrum of R6G was first measured was washed with distilled water. Then same R6G solution was loaded on the washed substrate and Raman spectrum was acquired. Signal intensities of four different peaks (612 cm^−1^, 773 cm^−1^, 1362 cm^−1^, and 1650 cm^−1^) from the bare substrate, sample load (1^st^), washed substrate, and sample load (2^nd^) were compared.

### Cell Culture

MCF7 cells (Korean Cell Line Bank, Korea), MDA-MB-231 cells (Korean Cell Line Bank, Korea), and SK-BR-3 cells (Korean Cell Line Bank, Korea) were cultured in Dulbecco’s modified Eagle’s medium (DMEM; Lonza, Switzerland) supplemented with 10 % (v/v) fetal bovine serum (FBS; Gibco, USA) and 1 % (v/v) antibiotic-antimycotic solution (A/A; Gibco, USA). MCF 10A cells (ATCC, USA) were cultured in mammary epithelial growth medium (MEGM; Lonza, Switzerland) supplemented with 100 ng/ml of cholera toxins (List Biological Laboratories, USA) and 1 % (v/v) A/A. All of the cells were cultured in a humidified incubator of 37 ℃ and 5 % CO_2_. Cells were detached and sub-cultured using 0.25 % trypsin-EDTA (Gibco, USA) when the cells reached 70-80 % confluency. For the preparation of the conditioned medium, the cell culture medium was aspirated and cells were washed with phosphate-buffered saline (PBS; Lonza, Switzerland) followed by cultured in an exosome-production medium for 48 hours. DMEM supplemented with 10 % (v/v) FBS which was centrifuged with 100,000 × g for 24 hours and 1% (v/v) A/A was used for exosome-production medium of MCF7 cells, MDA-MB-231 cells, and SK-BR-3 cells. MEGM was centrifuged with 100,000 × g for 24 hours and supplemented with 100 ng/ml of cholera toxins and 1 % (v/v) A/A for MCF 10A exosome-production medium.

### Exosome Isolation

Exosomes from each cell line were isolated from a conditioned medium based on differential ultracentrifugation (Optima™ XE-100, Beckman Coulter, USA) [57]. First, the conditioned medium was centrifuged at 2,000 × g for 10 minutes at 4 °C to remove live and dead cells. The supernatant without the pellet was transferred to fresh tubes and centrifuged 30 minutes at 10,000 × g, 4 °C to eliminate cellular debris. Then, the supernatant except the pellet was transferred to fresh tubes and centrifuged for 70 minutes at 110,000 × g, 4 °C. Followed by removing the supernatant, the pellet was resuspended by PBS and centrifuged with 110,000 × g for 70 minutes at 4 °C. At last, the PBS supernatant was removed completely and the pellet was resuspended by PBS or distilled water (DW; Daejung Chemicals & Metals Co., Korea). The purified exosome suspensions were stored at −80 °C in 10-μl aliquots for up to 1 year.

### Transmission Electron Microscopy (TEM)

Exosome suspension in PBS and 4% paraformaldehyde solution (PFA; biosesang, Korea) were mixed in a 1:1 volume ratio. Then, 5 μl of the mixed solution was dropped onto the 200 mesh formvar/carbon copper grid (Electron Microscopy Sciences, USA), allowing it to almost dry for 20 minutes. The grid was washed by depositing onto the DW drop with the grid membrane side down (5 minutes each, repeat 5 times). Followed by grid washing, negative staining was performed by depositing for 10 minutes with the gird membrane side down on a drop of 1% (w/v) uranyl acetate (Polysciences, USA). The remaining solution on the grid was removed using filter paper (Whatman, UK) and air-dried. Images are acquired at 200 kV using Talos F200X (FEI, USA). For AgNP analysis, AgNPs detached from the substrate using scalpel were suspended in DW and dropped on the 200 mesh formvar/carbon copper grid. The dried grid was then analyzed at 200 kV by Talos F200X.

### Nanoparticle Tracking Analysis (NTA)

Exosome suspension in PBS was diluted 1:100 in PBS. The diluted solution was analyzed for size distribution and concentration by NanoSight NS300 (Malvern Panalytical, UK) equipped with 532 nm laser. For consistency, detection threshold and camera level were set to 5 and 12 respectively. For each sample, three videos at 25 fps for 10 seconds were analyzed by NTA 3.2 software.

### Surface-Enhanced Raman Spectroscopy (SERS)

All the samples were prepared by dropping the solution on the AgNP substrate and left to dried condition. Raman spectra of samples were measured by dispersive Raman spectrometer (ARAMIS, Horiba Jobin Yvon, France). For R6G Raman spectra, the spectrometer equipped with a 514 nm He-Ne laser and ×50 long working distance lens was set to 300 μm of hole depth and 100 μm of slit size. The focused light with 0.313 mW laser power scanned the substrate for 5 s with 1 measurement number in the spectral range of 400 ∼ 1700 cm^−1^. For exosomal Raman spectra, a 633 nm Ar laser and ×50 long working distance lens were equipped with 300 μm of hole depth and 100 μm of slit size. 1.25 mW of focused laser scanned the substrate for 5 s with the measurement number of 1 in the spectral range of 300 ∼ 1800 cm^−1^.

### Data Analysis

Quantitative properties of the AgNP substrate (*i.e.* particle size distribution and covered area and count of particles) were acquired by Fiji software (https://imagej.net/Fiji). Surface images of the substrate were converted to binary images and “Analyze Particles” was executed to find particles with circularity between 0.8 and 1.0. Baseline correction and normalization of Raman spectra were conducted by BEADS toolbox in MATLAB (MathWorks, USA) (Figure S8, Supporting Information) [62]. The preprocessed spectra of exosomes were then analyzed by principal component analysis (PCA) using MATLAB. Dataset constituting a spectral range of 300 ∼ 1800 cm^−1^ was used as input data and calculated to acquire a covariance matrix. Eigenvectors from the covariance matrix were used as principal components in order of increasing magnitude for dimensional reduction of multidimensional data. Here, three eigenvectors were employed that reduce the variance of the entire data to a dimension that explains about 35 % of the variance. All of the data post-processed using MATLAB or OriginPro software (OriginLab, USA).

## Author Contributions

After the project was initiated by J. S. J. and M.Y.. H.N. and J.E.P. conducted experiments and analyzed data. J.S.H. and S.J.K. analyzed data. S.K. performed experiments. H.N. prepared the draft manuscript. †These authors contributed equally.

---

**Figure S1.**
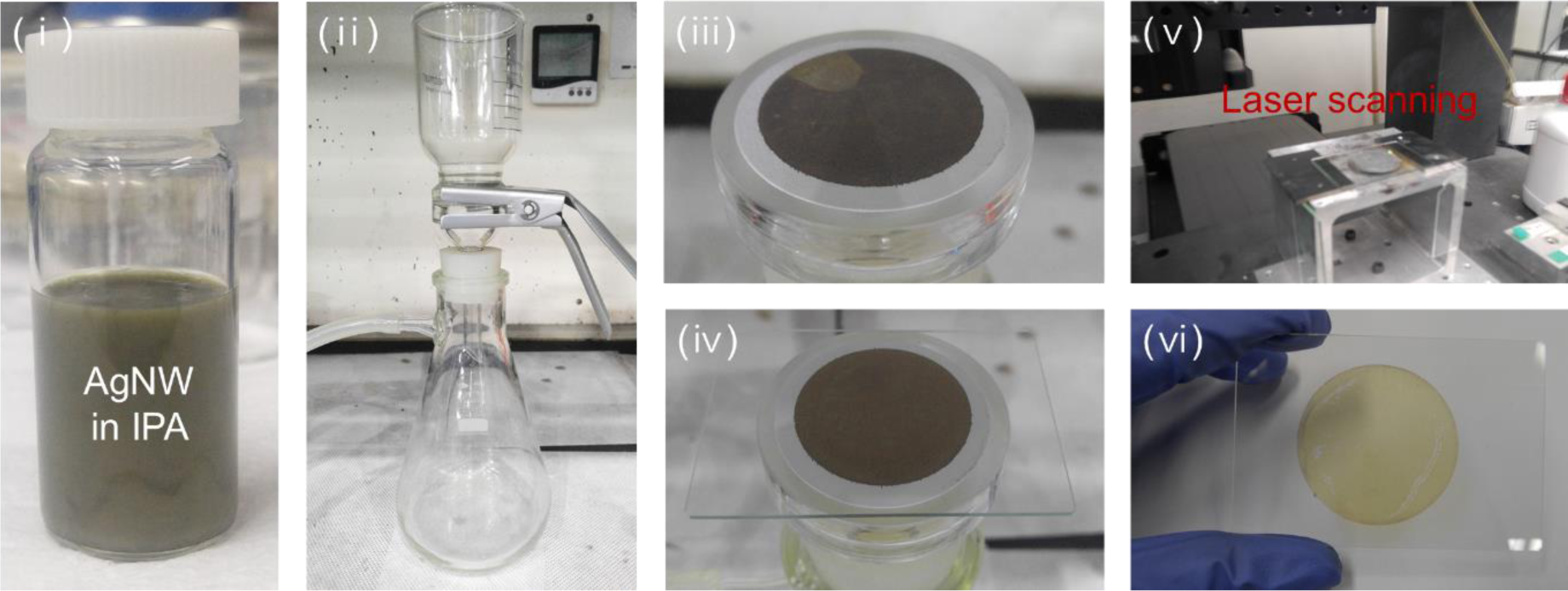
AgNP substrate manufacturing procedure. (ⅰ-ⅳ) AgNWs in AgNW solution were adhered to the glass substrate by vacuum filtration. (ⅴ, ⅵ) Laser ablation and melting-based fabrication of AgNP substrate from AgNW substrate.

**Figure S2.**
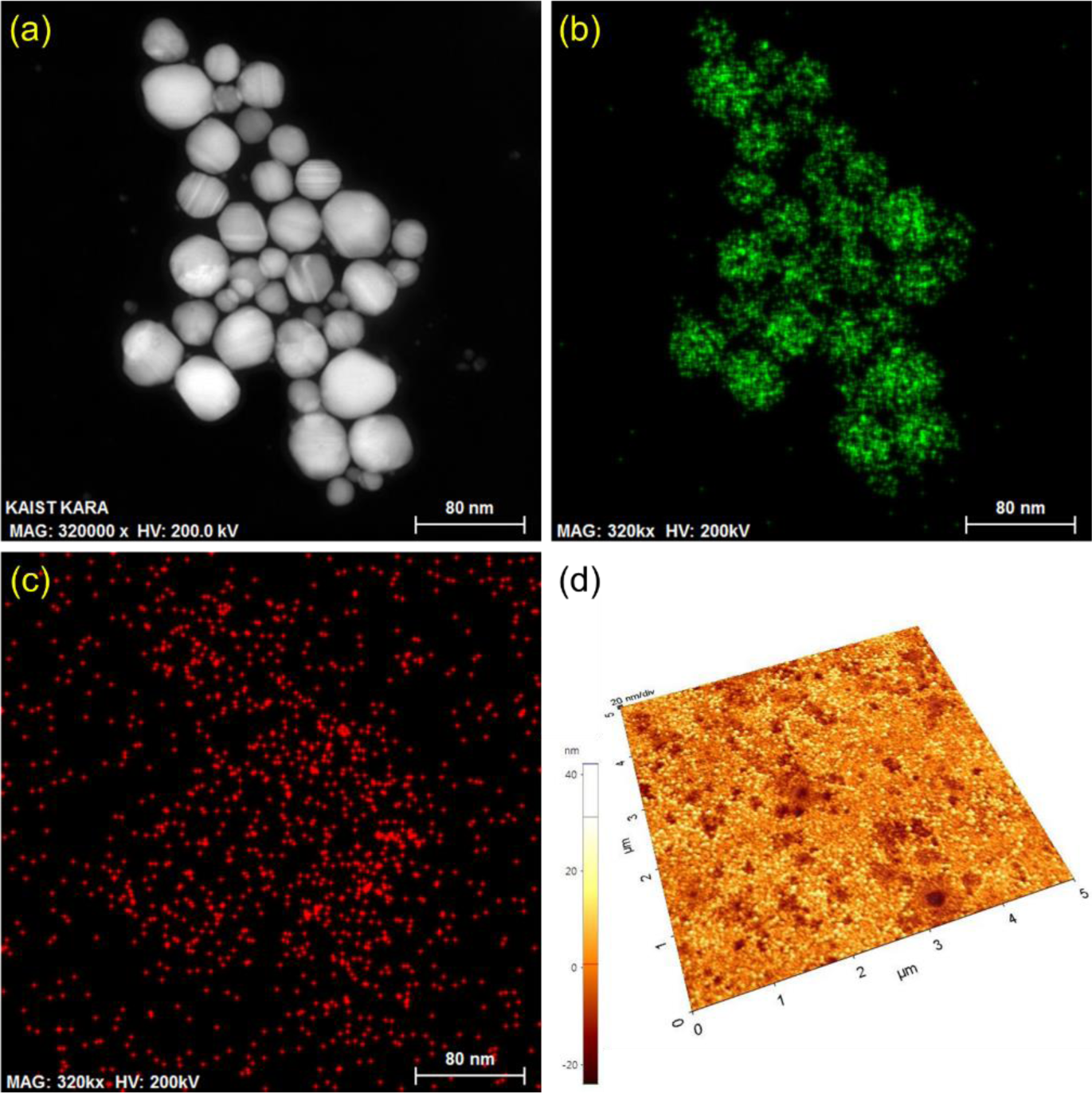
(a) Morphological analysis of AgNP substrate. (b-c) Energy dispersive spectrometer reveals the distribution of Ag (b) and Oxygen (c) on fabricated AgNPs. (d) AFM analysis of the surface of the AgNP substrate.

**Figure S3.**
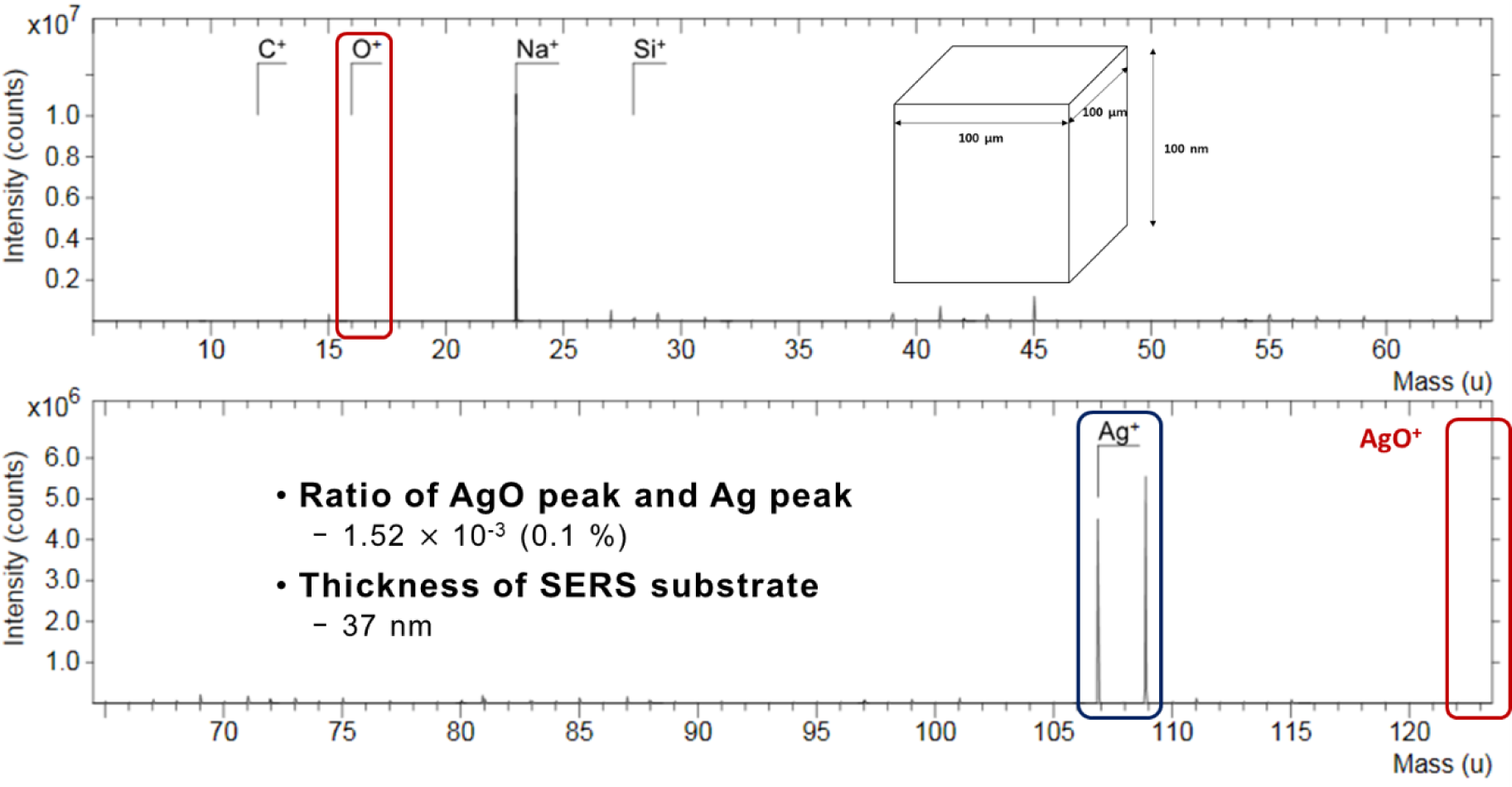
Surface profiling of AgNP substrate by TOF-SIMS.

**Figure S4.**
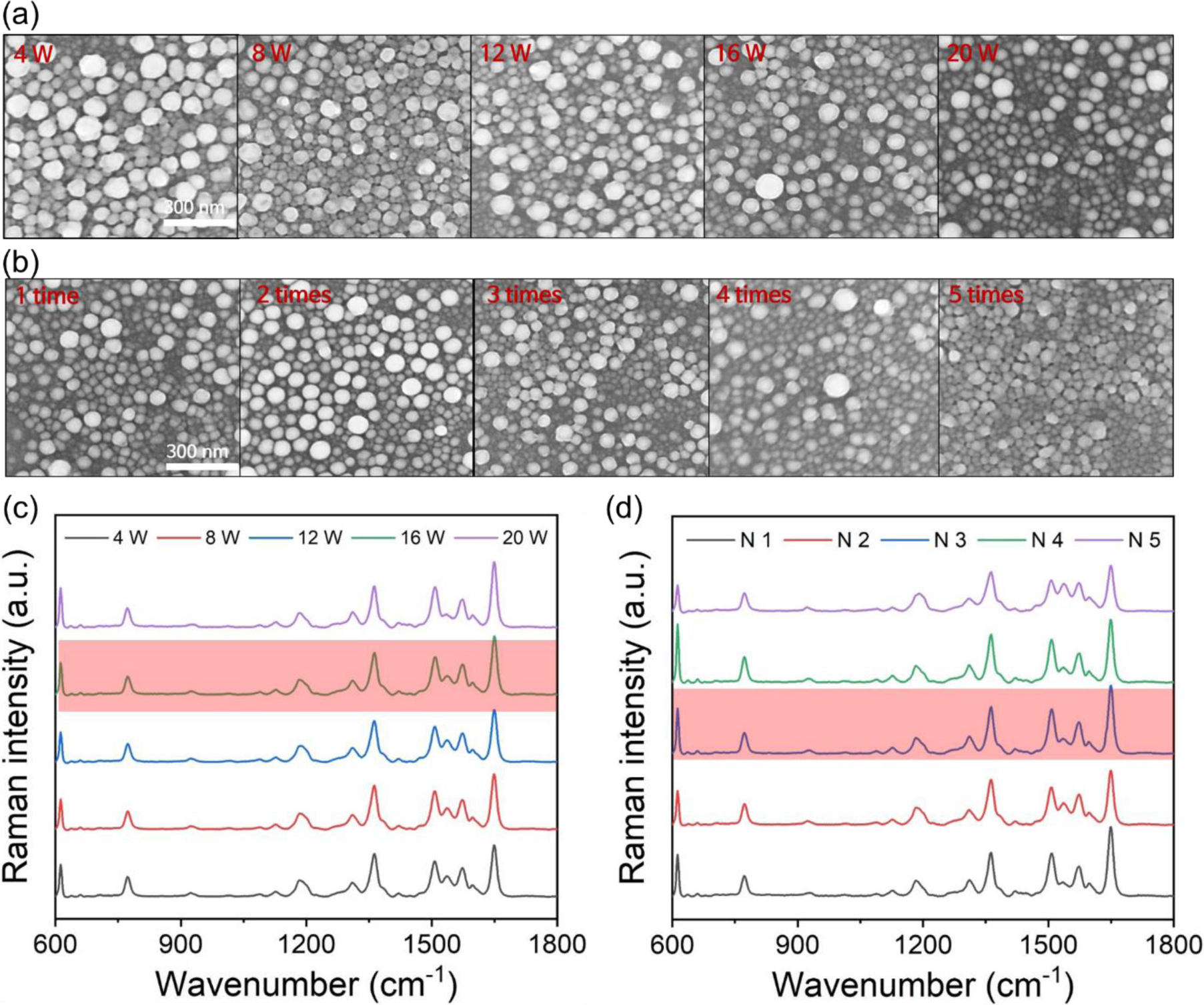
Functional evaluation under various laser conditions. (a) Surface morphology with increasing laser power. (b) Surface morphology by varying the laser ablation times. (c) Raman spectra of R6G under different laser powers. (d) Raman spectra of R6G under different laser ablation times.

**Figure S5.**
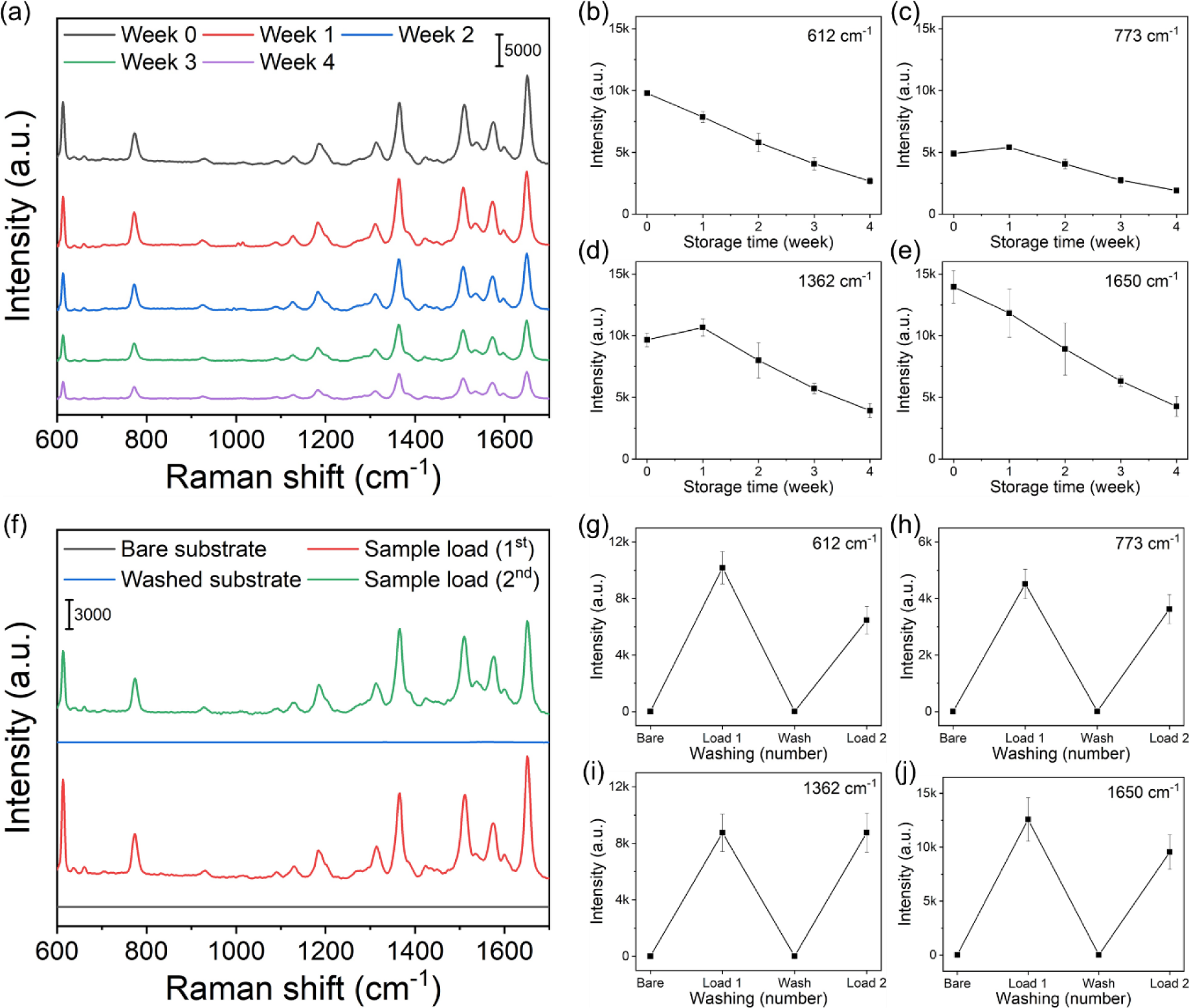
(a) Stability evaluated by obtaining Raman spectra of R6G every week. (b-e) Comparative analysis of specific wavenumbers for stability evaluation (error bar: standard deviation). (c) Renewability evaluation by subsequent sample loading after washing. (g-j) Signal intensities of four wavenumbers were compared for the estimation of renewability (error bar: standard deviation).

**Figure S6.**
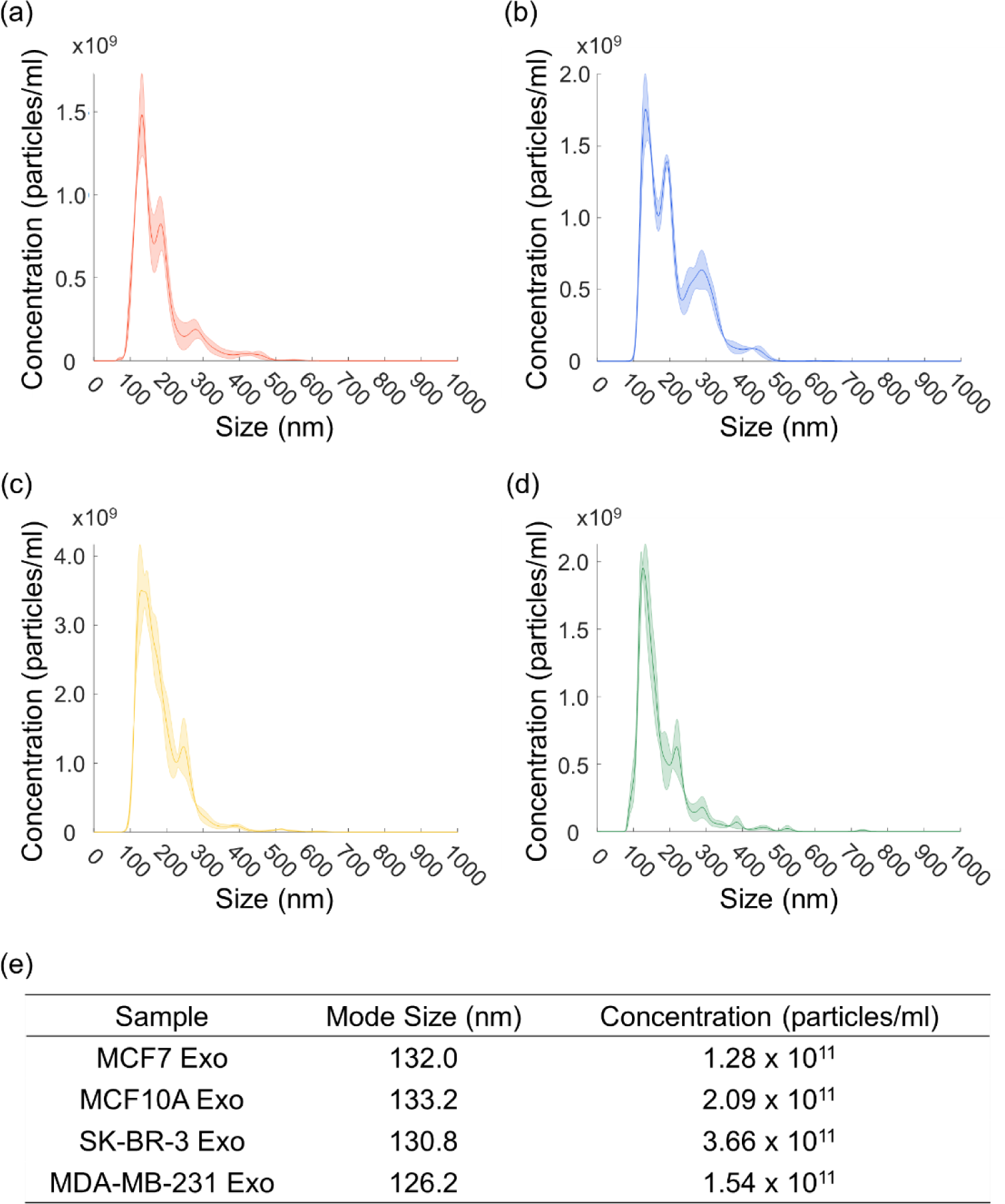
Quantitative analysis of exosome. Size distribution of four different cell line-derived exosomes were analyzed using NTA: (a) MCF7 exosome, (b) MCF 10A exosome, (c) SK-BR-3 exosome, (d) MDA-MB-231 exosome (shaded area: standard deviation). (e) Mode size and concentration of each isolated exosome suspension.

**Figure S7.**
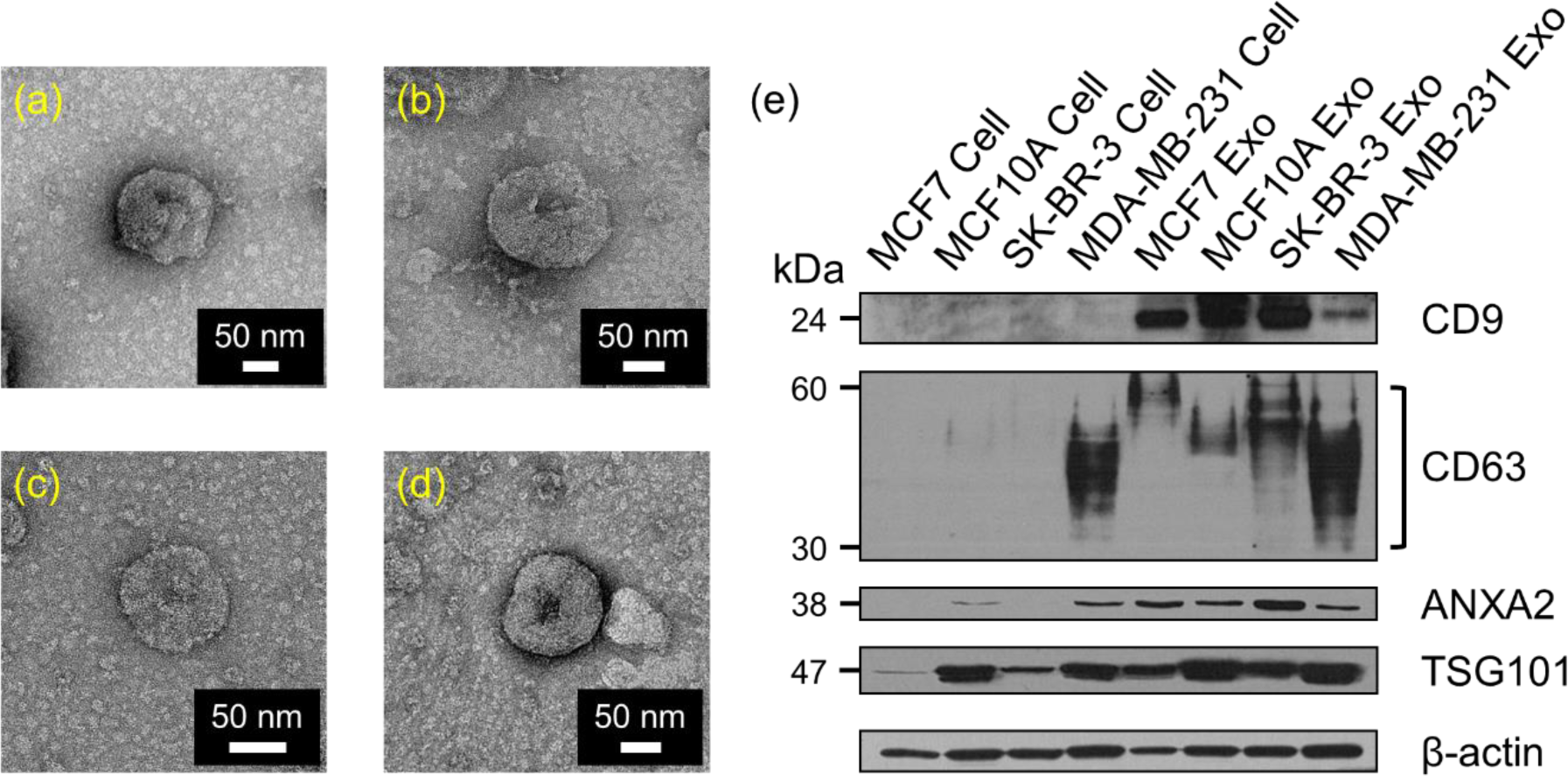
Qualitative analysis of exosome. Representative TEM images of each cell line-derived exosome: (a) MCF7 exosome, (b) MCF 10A exosome, (c) SK-BR-3 exosome, (d) MDA-MB-231 exosome. (e) Western blot of each cell line and cell line-derived exosome.

**Figure S8.**
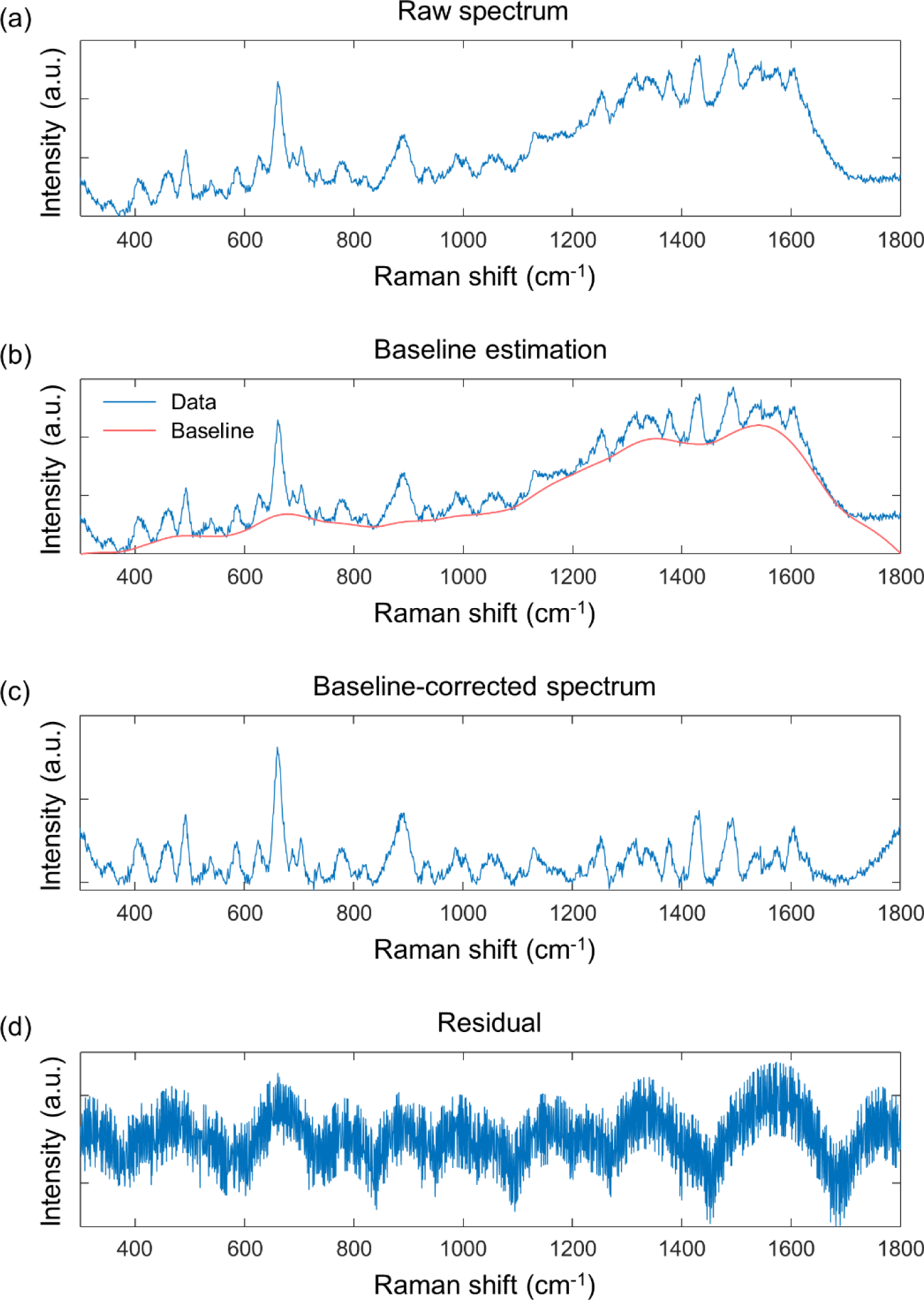
Raman spectra baseline correction by BEADS toolbox in MATLAB [1].

**Figure S9.**
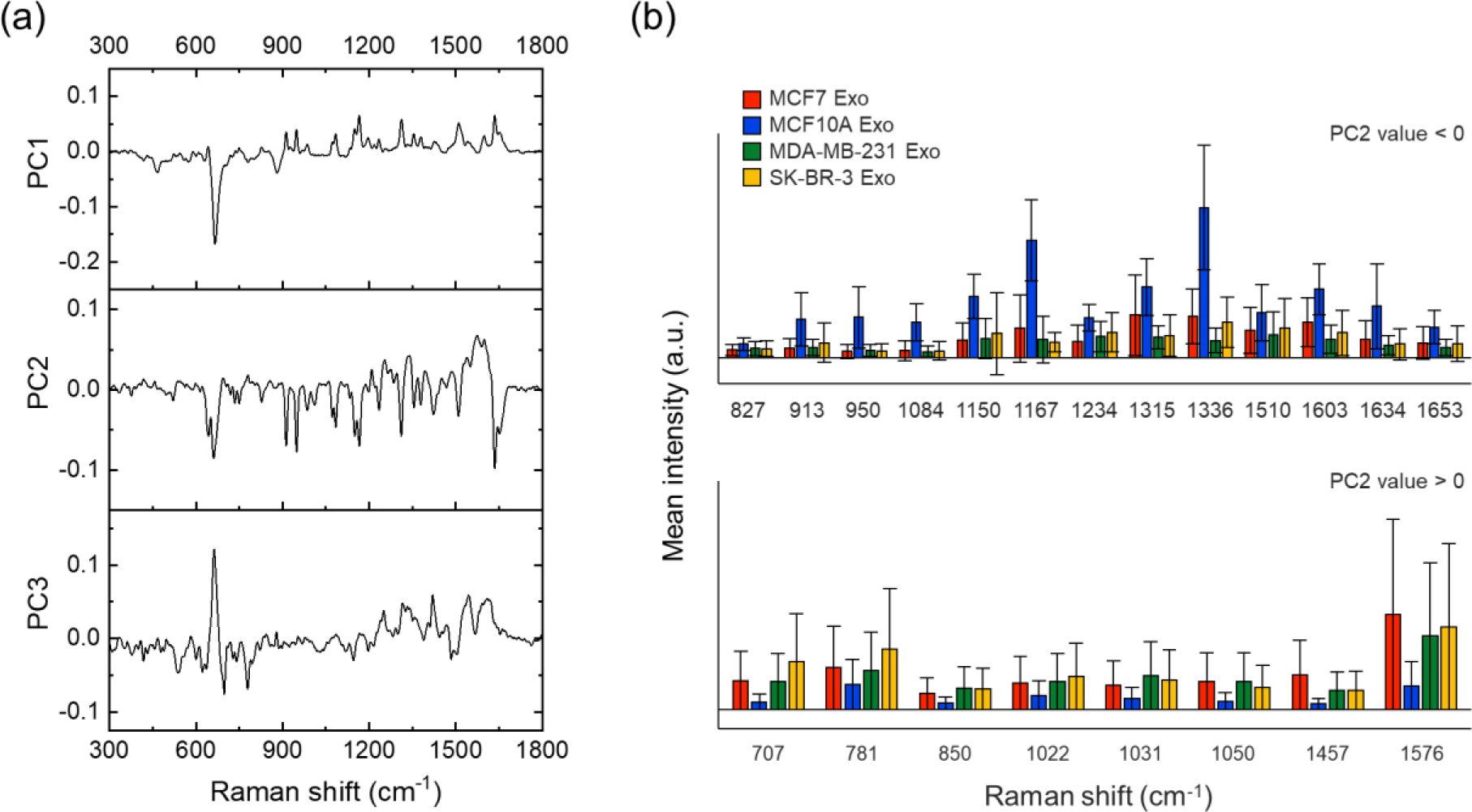
(a) Loading plots for each principal component. (b) Mean signal intensities of specific wavenumbers were compared based on PC2 (error bar: standard deviation)

**Figure S10.**
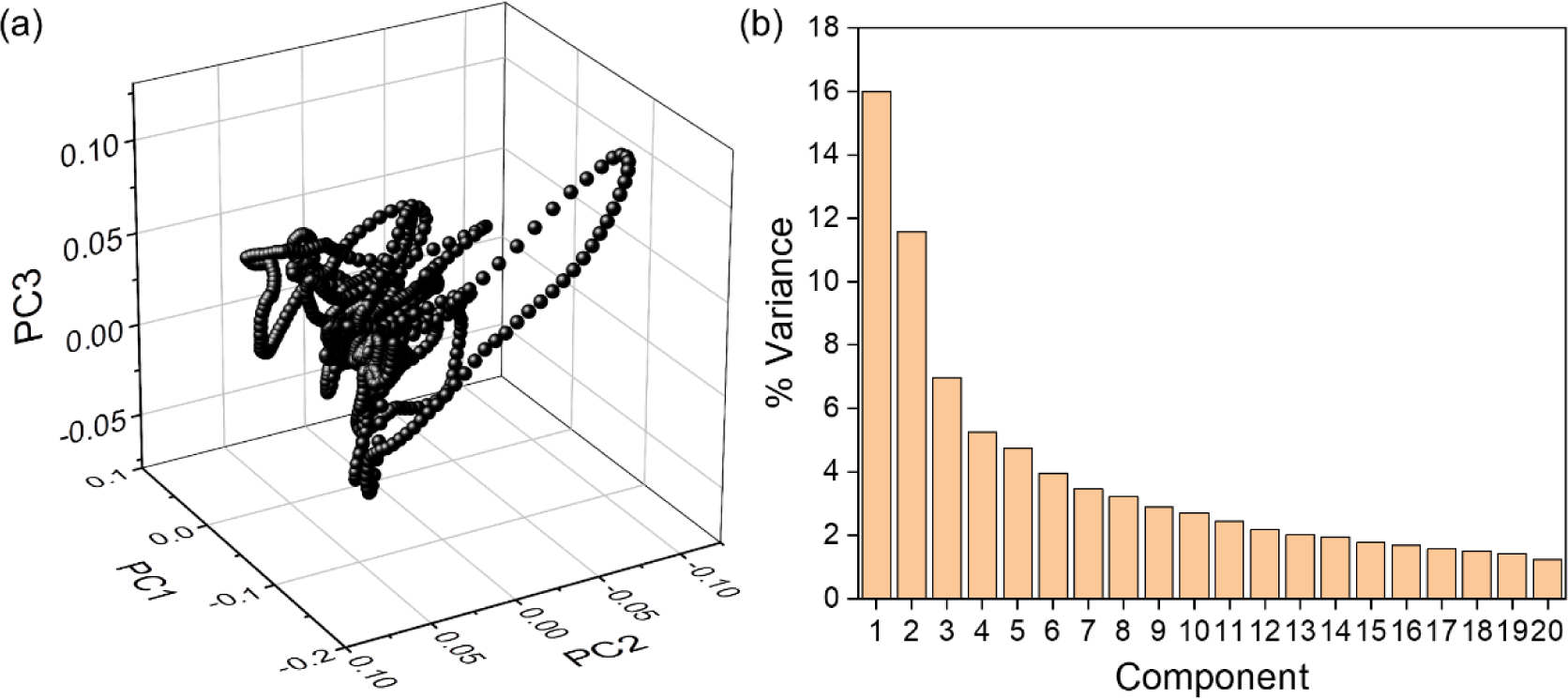
(a) Loading plot within the spectral range (300 ∼ 1800 cm^−1^). (b) Scree plot with respect to principal components.

**Table S1.**
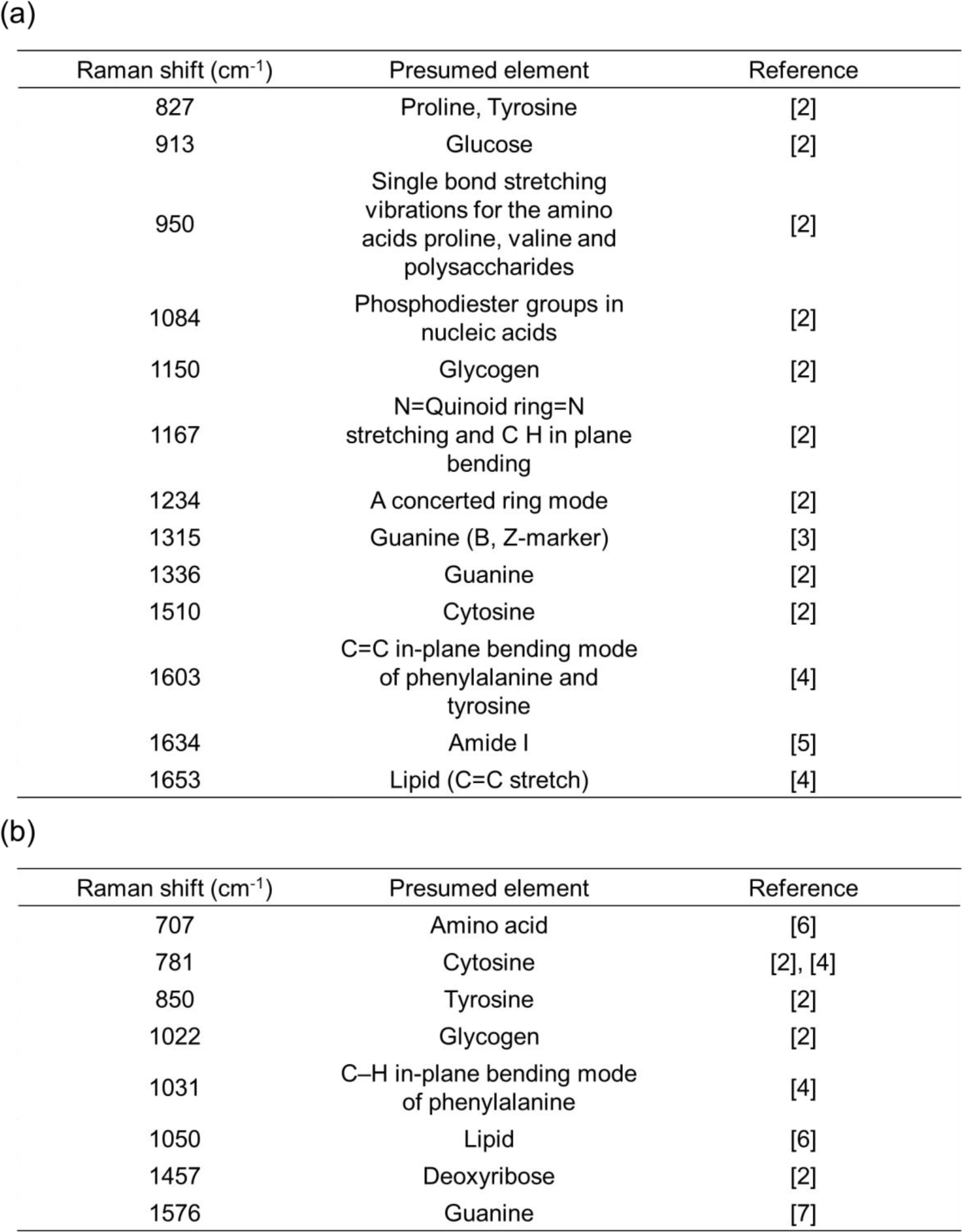
(a) Potential candidate substances for specific wavenumbers based on PC2 value < 0. (b) Potential candidate substances for specific wavenumbers based on PC2 > 0.

| Raman shift (cm<sup>-1</sup>) | Presumed element | Reference |
| --- | --- | --- |
| 827 | Proline, Tyrosine | [2] |
| 913 | Glucose | [2] |
| 950 | Single bond stretching vibrations for the amino acids proline, valine and polysaccharides | [2] |
| 1084 | Phosphodiester groups in nucleic acids | [2] |
| 1150 | Glycogen | [2] |
| 1167 | N=Quinoid ring=N stretching and C H in plane bending | [2] |
| 1234 | A concerted ring mode | [2] |
| 1315 | Guanine (B, Z-marker) | [3] |
| 1336 | Guanine | [2] |
| 1510 | Cytosine | [2] |
| 1603 | C=C in-plane bending mode of phenylalanine and tyrosine | [4] |
| 1634 | Amide I | [5] |
| 1653 | Lipid (C=C stretch) | [4] |

| Raman shift (cm<sup>-1</sup>) | Presumed element | Reference |
| --- | --- | --- |
| 707 | Amino acid | [6] |
| 781 | Cytosine | [2], [4] |
| 850 | Tyrosine | [2] |
| 1022 | Glycogen | [2] |
| 1031 | C-H in-plane bending mode of phenylalanine | [4] |
| 1050 | Lipid | [6] |
| 1457 | Deoxyribose | [2] |
| 1576 | Guanine | [7] |

